# Concurrent model evidence computation and posterior sampling in continuous attractor network subspaces

**DOI:** 10.64898/2026.09.02.748767

**Authors:** Yi Ren, Zimei Chen, Hayden Deans, Ying Nian Wu, Wen-Hao Zhang

## Abstract

Extensive studies suggest the brain performs Bayesian inference to infer the latent world states. It is a fundamental neuroscience question that how canonical recurrent neural circuits in the brain implement Bayesian inference. Many existing theoretical studies focused on how the recurrent circuits compute the posterior, while largely overlooking how the circuits compute the model evidence (normalization constant in Bayes’ theorem), a key quantity that measures how well a model explains the observed data. Thus, it remains largely unknown about how the recurrent circuits compute the model evidence. The present study performs rigorous theoretical analyses of the continuous attractor networks, a canonical recurrent circuit model, and reveals that the nonlinear circuit dynamics can simultaneously compute the posterior and model evidence in first two dominant subspaces within the circuit dynamics. Specifically, the circuit dynamics in the stimulus feature subspace implements the Langevin posterior sampling, and the circuit dynamics in the subspace of total neuronal activity computes the model evidence in a way analogous to the evidence lower bound in stochastic variational inference. We further extend the circuit model to compute the model evidence of multiple inputs, and simulations validate the computation in the network. Our work for the first time reveals the concurrent model evidence and posterior sampling in subspaces in continuous attractor networks, significantly deepen our understanding of the computational algorithms adopted by the neural circuits.

## 1 Introduction

The brain lives in a world of uncertainty and ambiguity, implying the observed inputs are often noisy and incomplete, necessitating the need of inference of true world states based on the observed inputs. Mathematically, this can be well described as the Bayesian inference. Supposing a latent world state *z* and the observed input *x*, the brain computes the posterior *p*(*z*|*x*) via the Bayes’ theorem (Eq. 1),

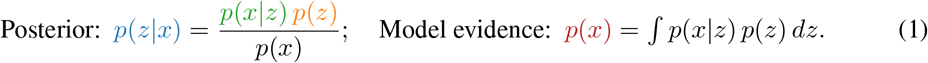

*p*(*x*|*z*) is the likelihood, *p*(*z*) is the prior. And *p*(*x*) is the model evidence (ME) measuring how well the model can explain the observation. Extensive cognitive studies have suggested that cognitive computations across domains aligns with the above Bayesian principles, giving rise to the influential concept of the “Bayesian brain” [1, 2]. These include visual processing [3], multi-sensory integration [4], decision-making [5, 6], causal inference [7], sensorimotor learning [8], etc.

The Bayesian computation at the cognitive level emerges from the neural circuits in the brain. It is a fundamental question in neuroscience to understand how the recurrent neural circuits implement Bayesian inference. Many theoretical studies have investigated the posterior *p*(*z*|*x*) computation in recurrent neural circuits, and proposed various potential neural circuit algorithms, which can be classified into two categories including the deterministic algorithms based on probabilistic population codes [2, 5, 9–12], and stochastic algorithms based on Bayesian sampling [13–22].

Despite the large body of studies on posterior computation in neural circuits, however, previous studies have largely overlooked how neural circuits compute the ME *p*(*x*) in either neuroscience and machine learning (ML) research (Eq. 1). To our best knowledge, there is only one related study (see Discussion) [10], and this problem has been basically untouched in the past 15 years. Although *p*(*x*) is not important for posterior *p*(*z x*) computation, it is critical for Bayesian model learning and comparison through telling how well the model as a whole can explain the input *x* by integrating all latent states *z*. In **neuroscience**, the ME is the key quantity underlying essential cognitive functions such as causal inference [7, 23, 24], decision-making [5, 25, 26], and learning and synaptic plasticity [27, 28], etc. However, the neuroscience field has a large knowledge gap in understanding ME computation in recurrent neural circuits, impeding our understanding of circuit algorithms underlying those cognitive functions. Meanwhile, in **machine learning**, the ME acts as the objective function in generative models. Although many efficient algorithms have been proposed to estimate the ME, e.g., variational autoencoders [29] and diffusion models [30, 31], to our best knowledge, none of them studied how these algorithms can be embedded into neural networks. Rather, neural networks are just used as encoders/decoders to connect input spaces with latent spaces. If we can pinpoint the ME calculation in recurrent circuits, the circuits with automatic ME estimation may bridge the gap between neural networks and generative models, and inspire the next-generation AI systems.

To provide theoretical and algorithmic understanding of how biologically plausible recurrent neural circuits compute the model evidence during the posterior computation, the present study considers a canonical recurrent circuit model – continuous attractor networks (CANs) that consists of excitatory neurons and a pool of global inhibitory neurons [32–35]. The circuit model receives feedforward inputs conveying the observed stimulus feature, and top-down inputs conveying the prior about the stimulus feature (Fig. 1A). After rigorous theoretical analyses of the nonlinear CAN dynamics, we analytically derive the dynamics on its first two dominant subspaces: the **stimulus feature subspace** and the **total neuronal activity** subspace, respectively. Then we surprisingly find the first two dominant subspaces concurrently compute the posterior and the model evidence, respectively.

**Figure 1:**
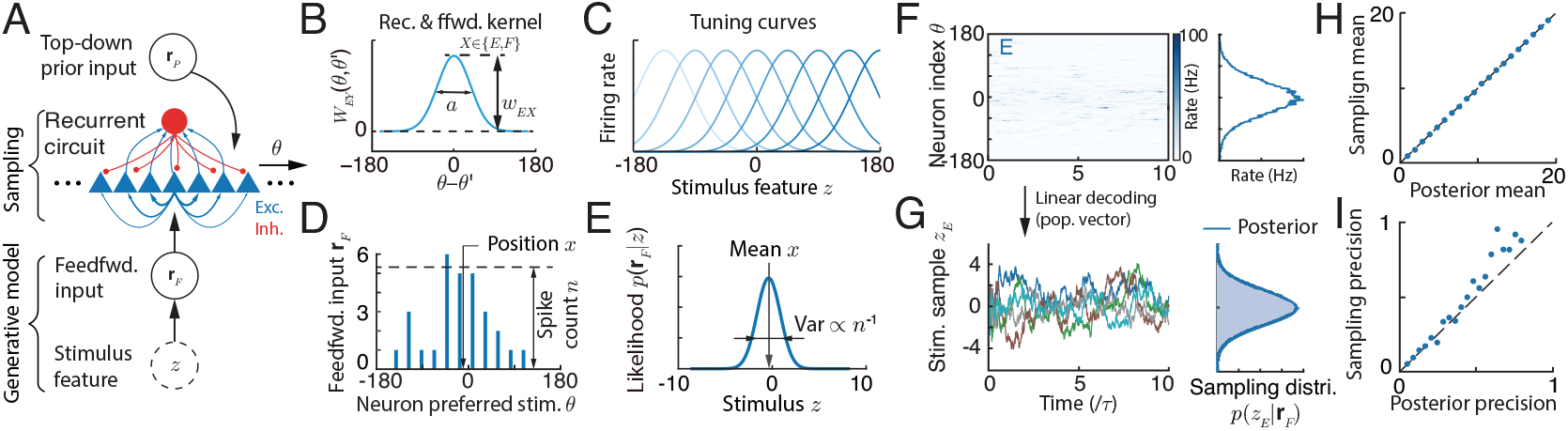
The recurrent circuit model. (A) Circuit model architecture. (B) Recurrent connection kernel. (C) E neurons’ tuning curves (mean firing rates) to the stimilus feature *z*. (D) A snapshot of the feedforward sensory input **r**_*F*_ . (E) The stimulus likelihood given the feedforward input. (F) The spatiotemporal E neuronal activity. Right: time-averaged E activity. (G) The stimulus feature samples over time, which is read out as the E activity’s position in the stimulus feature space. Right: The histogram of the stimulus feature samples over time, which converges to the posterior. (H-I) The circuit with fixed synaptic weights flexibly sample posteriors with different means and precisions. The posterior parameter is manipulated by changing the intensity and location of feedforward inputs.

Specifically, the circuit dynamics in the stimulus feature subspace implements the Langevin posterior sampling (Sec. 4), while the total activity subspace dynamics computes the logarithm of (unnormalized) model evidence (Sec. 5). Further analysis on the non-equilibrium circuit subspace dynamics reveals that the circuit utilizes its **multiscale temporal** subspace dynamics to compute the model evidence in a way analogous to the evidence lower bound (ELBO) in stochastic variational inference: the circuit first utilizes the relatively **fast** stimulus feature subspace dynamics to sample the posterior, and then the circuit utilizes the relatively **slow** dynamics in the total activity subspace to compute the model evidence by integrating over the posterior samples from the stimulus feature subspace (Fig. 3). We also extend the circuits to compute the model evidence of multiple inputs (Fig. 4), and the simulations confirm the circuit can compute the model evidence sufficiently well.

### Significance and contributions

**1)** The **first** algorithmic understanding about how the recurrent circuits concurrently calculates model evidence and posterior sampling in its subspaces, deepening our understanding of circuit’s Bayesian computations. **2)** We show the non-equilibrium circuit dynamics is analogous to stochastic variational inference. **3)** Provide a precise map between nonlinear circuit dynamics and modern Bayesian algorithms. **4)** To ML: The recurrent circuit model automatically embeds modern Bayesian algorithms, potentially inspiring a new building block for future AI systems.

## 2. Continuous attractor networks: A canonical recurrent circuit model

The present study considers a canonical recurrent circuit model – continuous attractor networks (CANs) that consists of excitatory and a pool of inhibitory neurons (Fig. 1A, details in Appendix B). The CANs can be regarded as a cortical hypercolumn, and have been widely used in neuroscience research [18, 22, 33, 35, 36], including the primary visual cortex [36, 37], prefontal cortex [38, 39], the head direction system [32], place cells [40], and grid cells [41, 42], etc.

### E neurons’ dynamics

The E neurons in the circuit are selective for the 1D stimulus feature *z* in Eq. (1), e.g., orientation. Denote *θ*_*j*_ as the preferred stimulus feature of the *j*-th E neuron, and the preferred feature of all *N*_*E*_ E neurons, 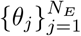 uniformly cover the whole feature space *z*, a setting widely adopted in neural circuit modeling. To facilitate math analysis, we change the summation of recurrent inputs over the neural space into continuous integration (Appendix B.1),

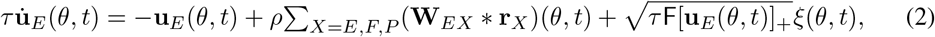

where **u**_*E*_(*θ, t*) and **r**_*E*_(*θ, t*) represent, respectively, the synaptic inputs and firing rates of neurons preferring *z* = *θ. X* denotes neuronal types with *E, F*, and *P* representing E neurons, sensory feedforward inputs and top-down prior inputs, respectively. *τ* is the time constant, *ρ* = *N/*2*π* is the neuronal density covering the stimulus feature space, and [*x*]_+_ = max(*x*, 0).

### Recurrent connection kernel

**W**_*Y X*_ (*θ*) is the recurrent connection kernel from neurons with type *X* to those with type *Y*, which are modeled as Gaussian functions in the model (Fig. 1B),

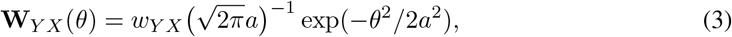

where the scalar *w*_*Y X*_ is the peak recurrent weight strength and is the only circuit parameter to be adjusted to realize Bayesian computation. *a* is the connection width across the stimulus feature space. The symbol ∗ denotes the convolution, i.e., **W**(*θ*) ∗ **r**(*θ*) = ∫**W**(*θ* − *θ*^*′*^)**r**(*θ*^*′*^)*dθ*^*′*^.

### Divisive normalization as the activation function

There is a pool of global inhibitory neurons that are driven by E neurons and provide global, divisive inhibition to E neurons. For simplicity, we incorporate their effects as the divisive normalization (DN) acting as the activation function of E neurons, rather than explicitly modeling their dynamics,

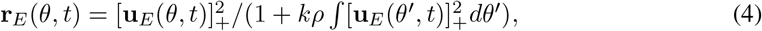

*k* (a positive scalar) characterizes the global inhibition strength. The divisive normalization is a canonical operation found in the cortex [43–45]. Note that the DN only acts as an activation function transferring the instantaneous synaptic input **u**_*E*_ to the firing rate **r**_*E*_, rather than normalizing the neuronal activity into a probability distribution with integration equal to 1.

### Feedforward sensory neural input (encoding observed feature *x*)

The feedforward sensory neural input **r**_*F*_ (*θ, t*) (Eq. 2) is stochastically evoked by the latent stimulus feature *z*, and is modeled as conditionally independent Poisson spikes with Gaussian tuning given a stimulus *z* (Fig. 1D), consistent with a large body of previous models (e.g., [9, 10, 18, 19]).

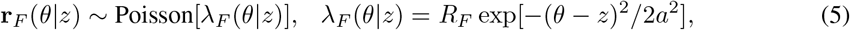

where *λ*_*F*_ (*θ*|*z*) is the mean firing rate. In simulating the rate-based circuit dynamics, **r**_*F*_ is approximated as a continuous Gaussian random variable with multiplicative noise.

### Top-down prior neural input

The neural circuit also receives a top-down neural input **r**_*P*_ that conveys the prior information about the stimulus feature *z* (Fig. 1A). For simplicity, the top-down prior input **r**_*P*_ has the similar form with the feedforward input but with different values of *R*_*F*_ and *z* in Eq. (5) (details in Appendix C6). The top-down prior input **r**_*P*_ is fixed over time and trials, reflecting the stable prior knowledge stored in the circuit. In contrast, the feedforward input **r**_*F*_ is stochastically generated over trials and over time, reflecting the noisy sensory transmission.

## 3 The generative model embedded in the circuit model

### Stimulus feature likelihood

The stochastic generation of the feedforward input **r**_*F*_ from the stimulus feature *z* (Eq. 5) is regarded as the generative process (Fig. 1A). The stimulus likelihood given a feedforward input **r**_*F*_, *p*(**r**_*F*_ |*z*) is derived as a Gaussian likelihood (Appendix C),

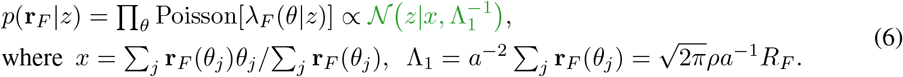

The likelihood mean *x* is regarded as the **observed stimulus feature**. The mean *x* and the precision Λ_1_ can be read out from **r**_*F*_ via a linear decoder called population vector [46, 47], which geometrically refer to **r**_*F*_ ‘s location in the stimulus feature space and the height (magnitude) of the feedforward input (Fig. 1E), respectively. Note that the Gaussian stimulus likelihood results from the Gaussian tuning of feedforward input *λ*_*F*_ (*θ*|*z*) (Eq. 5) [2, 9], rather than directly specified by hand.

### Stimulus feature prior

Similarly, the stimulus feature prior *p*(*z*) is represented by the top-down prior input **r**_*P*_ in the circuit, and its mean and precision can be read out from **r**_*P*_ via the population vector decoder as well (replacing the **r**_*F*_ by **r**_*P*_ in Eq. 6).

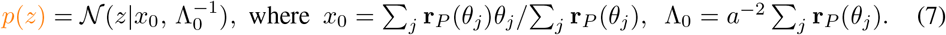

### Stimulus feature posterior

The posterior can be computed as a Gaussian distribution,

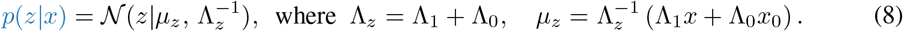

The posterior mean *µ*_*z*_ is a precision-weighted combination of the *x* and the prior mean *x*_0_: a more reliable observation (large Λ_1_) pulls *µ*_*z*_ toward *x*, and otherwise toward *x*_0_.

### Model evidence of the observed feature *x*

It can be analytically calculated as,

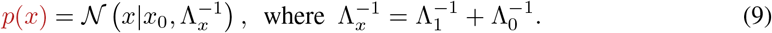

Note that we will study how the circuit calculates the model evidence of the observed feature *p*(*x*), rather than the evidence of the observed high-dimensional feedforward input *p*(**r**_*F*_). Here the *x* corresponds to the summary statistics of **r**_*F*_ in determining stimulus *z* likelihood (Eq. 6). Moreover, during the posterior computation (Eq. 1), the *p*(*x*) is a *scalar value* of probability evaluated at the observed feature *x*, instead of the whole distribution over all possible *x*. Comparing the precision of the posterior and the model evidence, the posterior’s precision Λ_*z*_ is the sum of the likelihood and prior precisions (Eq. 8), whereas the model evidence’s precision Λ_*x*_ is the *harmonic mean* of the likelihood and prior precisions, i.e., the sum of inverse precisions (Eq. 9). It remains largely unknown how circuits compute the harmonic mean of precisions involved in model evidence.

The rest of the paper will focus on how the nonlinear circuits can utilize its dynamics to automatically and concurrently compute the posterior *p*(*z*|*x*) and model evidence *p*(*x*). The analytical solutions of the Gaussian model provides a ground truth to validate the posterior and model evidence computation in the circuits. Despite the simplicity of the Gaussian model, we emphasize that it is still unknown about how recurrent circuits compute the posterior and model evidence even in this simple case.

### 3.1 Theoretical analysis of the circuit dynamics

To identify how the circuit can simultaneously compute the posterior and model evidence, we perform rigorous theoretical analyses of the circuit dynamics. We perform perturbative analysis of the nonlinear circuit dynamics, analytically identify the dominant modes (eigenvectors) of the perturbed dynamics, and then project the dynamics onto these modes. We eventually find the circuit dynamics performs posterior and model evidence computation in two dominant modes simultaneously.

#### Attractors

Given the feedforward and top-down inputs (Eq. 5), the synaptic input **u**_*E*_(*θ*) and firing rate **r**_*E*_(*θ*) of E neurons both have Gaussian bump attractor states (Fig. 1E-F; Appendix B.2),

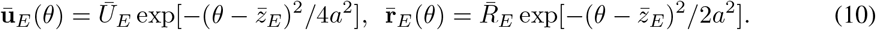

The Gaussian bump is located at 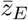 on the stimulus feature manifold, which is a weighted average of the observed feature *x* and the prior mean *x*_0_. *Ū*_*E*_ and 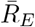 are the “height” of the population synaptic input and firing rate respectively, with solutions in Eqs. (12a-12b).

#### Perturbative analysis for dominant subspaces

The sensory noises and the internal Poisson variability perturb instantaneous neural responses from the attractor state (Eq. 10), i.e., **u**_*E*_(*θ, t*) = **Ū**_*E*_(*θ*) + *δ***u**_*E*_(*θ, t*). We then analytically derive the relaxation dynamics of the perturbation *δ***u**_*X*_ (*θ, t*) and its eigenvectors (Appendix B.3, [48]). The first two dominant eigenvectors correspond to the changes of position *z*_*E*_ and height *U*_*E*_ of the population activity respectively [33, 48],

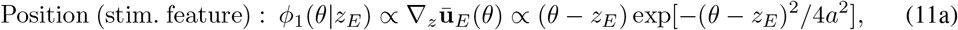

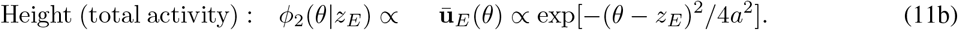

#### Dimensionality reduction for Bayesian computation in subspaces

We further project the neural dynamics (Eq. 2) onto the above two dominant eigenvectors, where the projection is computing the inner product between the neural dynamics and the eigenvector, i.e., ⟨*ϕ*(*θ*), *f* (*θ*) ⟩ = ∫*ϕ*(*θ*)*f* (*θ*)*dθ*, with *f* (*θ*) representing each term in Eq. (2). The projection yields the governing dynamics of the neurons on the stimulus feature subspace and height subspace (Appendix B.3), which is a critical step to explicitly identify the Bayesian computations embedded in the nonlinear circuit dynamics,

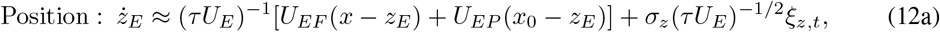

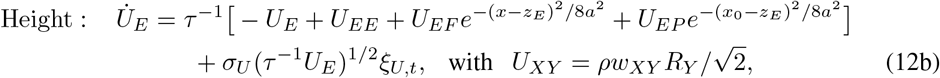

where *U*_*XY*_ is the population input magnitude from population *Y* to 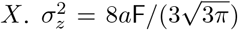 and 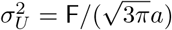 are all constants that don’t change with network activities. The approximation in Eq. (12a) comes from omitting negligible nonlinear terms in the circuit dynamics (Appendix B.3). *ξ*_*z,t*_ and *ξ*_*U,t*_ are independent Gaussian white noises in the position and height subspace respectively. Although both of them come from the internal Poisson variability (Eq. 2), they are independent because of the orthogonality of the position and height eigenvectors (Eqs. 11a-11b).

There are several important structures about the two subspace dynamics.

1. The feedforward and prior **input magnitude** *U*_*EF*_ and *U*_*EP*_ are proportional to the likelihood and prior **precision** respectively, e.g., *U*_*EF*_ ∝ *R*_*F*_ ∝ Λ_1_ (Eq. 6), and similarly for *U*_*EP*_ ∝ Λ_0_. This will be used to align the position dynamics with the **Langevin posterior sampling** (Sec. 4).
2. The height dynamics is modulated by the instantaneous position *z*_*E*_ (Eq. 12b, Gaussian terms), even if the position and height eigenvectors are orthogonal (Eqs. 11a-11b). This is because the height eigenvector is dependent on the instantaneous *z*_*E*_ (Eq. 11b) due to the nonlinear circuit dynamics. This modulation is crucial for the **model evidence** computation (Sec. 5).

## 4 Bump position subspace for Langevin posterior sampling

A recent study suggested the E bump position *z*_*E*_ (Eq. 12b, Fig. 1G) implement Langevin sampling of the stimulus posterior *p*(*z*|*x*) [22]. We briefly explain the mechanism below (details in Appendix B). To facilitate the understanding, we list the Langevin posterior *p*(*z*|*x*) sampling dynamics (Eq. 1),

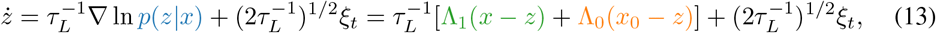

with *τ*_*L*_ *>* 0 being the sampling time constant. Meanwhile, Eq. (12b) can be converted into a form similar to the Langevin sampling (Eq. 13) by expressing *U*_*XY*_ as precisions Λ of likelihood and prior (Eq. 6). Combining *U*_*XY*_ definition (Eq. 12b) and the likelihood and prior precisions (Eqs. 6, 7),

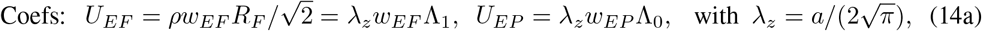

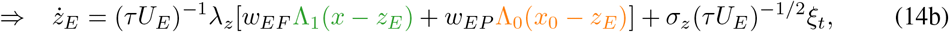

Comparing Eqs. (14b) and (13), realizing Langevin posterior sampling in the bump position subspace only requires the ratio between the drift and diffusion coefficients be the same as the one in Langevin sampling (Eq. 13), through setting the feedforward and prior input weights,

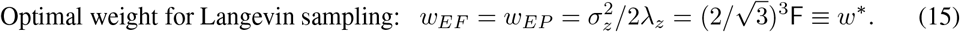

Simulation results confirm our theoretical finding. Moreover, once the weight *w*_*EF*_ = *w*_*EP*_ is set at the optimal value, the circuit with fixed synaptic weights can flexibly sample posteriors with different parameters (Fig. 1H-I), because *λ*_*z*_ and *σ*_*z*_ are both unvaried with network activities. Here, we manipulate the location and strength of the prior and feedforward inputs (Eq. 5) to change posteriors.

## 5 Bump height subspace computes the logarithm of model evidence

The model evidence *p*(*x*) decreases with the disparity between the observed feature *x* and the prior mean *x*_0_, |*x* − *x*_0_| (Eq. 9). Meanwhile, the time-averaged value of the E bump height *Ū*_*E*_ also decreases with |*x* − *x*_0_| (Fig. 2B), due to the nonlinear Gaussian terms in Eq. (12b). This can be intuitively understood by a simple example that the sum of two Gaussian-profile inputs will become smaller if their peaks are further apart (Fig. 2A). The suppression of *Ū*_*E*_ by feature disparity |*x* − *x*_0_| was widely observed in experiments on single neuron responses, such as the cross-orientation suppression in V1 [44, 49, 50], and the surround suppression in MT [51–53], and the decision-making circuits in LIP [26, 54–56]. These suppression on single neurons is also reproduced by our circuit (Fig. A1), suggesting its biological plausibility. Then we analyze the bump height *U*_*E*_ dynamics (Eq. 12b) to find how it can potentially compute the model evidence *p*(*x*).

**Figure 2:**
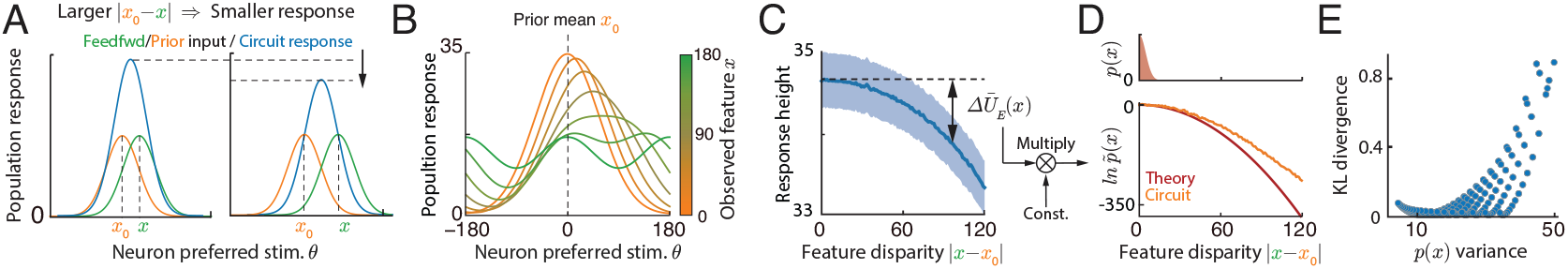
The total activity (bump height) of E neurons computes the model evidence *p*(*x*). (A) The schematic illustrates the mechanism of the model evidence computation in the circuits. When changing the disparity between the observed feature *x* (from feedforward input) and the prior mean *x*_0_ (from the top-down prior input), the magnitude (height) of the E population responses decreases with the disparity |*x* −*x*_0_|, analogous to the decrement of the model evidence *p*(*x*) (Eq. 9). (B) The time-averaged E population responses with various observed feature *x*, during which the prior mean *x*_0_ is fixed at 0. (C) The height of E population responses decreases with the disparity |*x* − *x*_0_|. Shaded region: 1 std. (D) The logarithm of (unnormalized) model evidence ln 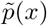 can be decoded by multiplying the decrement of E bump height with a constant (Eq. 18). Top: the ground truth model evidence *p*(*x*). (E) The KL divergence from the ground truth *p*(*x*) and the one computed by the circuits. Each dot is a result under a combination of likelihood and prior precisions. The circuit’s computed model evidence will have larger error with increasing variance of model evidence.

### 5.1 Mean bump height in the equilibrium state

We calculate the mean bump height *Ū*_*E*_ (*x*) in response to the observed feature *x* (the prior mean is fixed at *x*_0_). It is worth noting that finding *Ū*_*E*_ (*x*) requires averaging over **two sources of variability** (Eq. 12b): **One** is its own variability *ξ*_*U,t*_ with average of zero (Eq. 12b). **Another** variability is the fluctuation of *z*_*E*_ due to stimulus posterior sampling (Eq. 12a), whose average corresponds to finding the expectation of the Gaussian terms in Eq. (12b) over the posterior of *z*_*E*_, i.e., 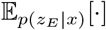,

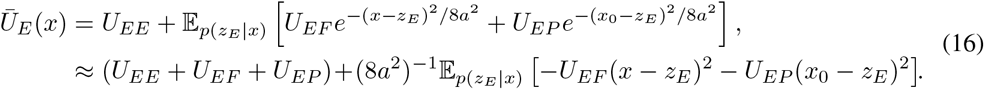

To directly connect *Ū*_*E*_ (*x*) with the logarithm of likelihood and prior, Taylor expanding the Gaussian terms to the first order, e.g., 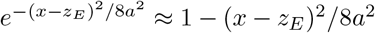, which works well since the 8*a*^2^ is large, i.e., E neurons are broadly connected in the stimulus feature space (Eq. 3, Fig. 1B-C). Later we will use simulation to evaluate the validity of the expansion (Fig. 2D). In addition, utilizing the input magnitude *U*_*EF*_ and *U*_*EP*_ are proportional to the likelihood and prior precision, respectively (Eq. 14a), the quadratic terms are proportional to the logarithm of likelihood and prior (Appendix E.1),

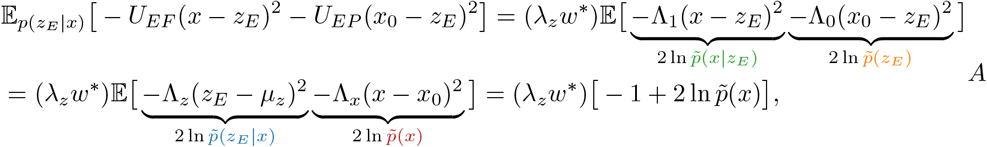

where Λ_*z*_(*z*_*E*_ − *µ*_*z*_)^2^ ∼ *χ*^2^(1) satisfies a chi-square distribution with one deg_J_ree of freedom and has a mean value of 1. 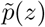 is the unnormalized distribution with 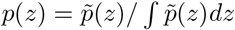, and similarly For 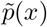. Eventually, the mean of bump height is found as,

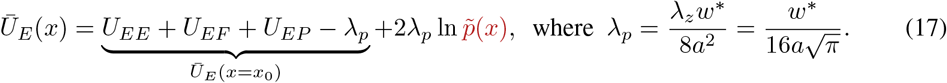

Importantly, the sum of the first four RHS terms in Eq. (17) is the mean bump height *Ū*_*E*_ (*x*_0_) when *x* = *x*_0_ (the prior mean). Hence, the **decrement of the mean bump height**, Δ*Ū*_*E*_ (*x*) ≡ *Ū*_*E*_ (*x*) − *Ū*_*E*_ (*x*_0_), results from disparity between |*x* − *x*_0_| is proportional to the logarithm of model evidence ln 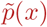 (unnormalized up to a constant), i.e.,

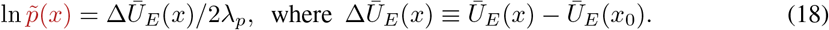

We simulate the network to verify the above theoretical analysis. The circuit model’s estimation ln 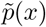, read out by using Eq. (18) (Fig. 2C), aligns with the ground truth when the observed feature *x* is close to the prior mean *x*_0_, while it deviates when *x* is far from *x*_0_ with upward bias (Fig. 2D). This is understandable because our theoretical analysis of Δ*Ū*_*E*_ (*x*) (Eq. 16) is based on the first-order Taylor expansion of the nonlinear Gaussian terms in Eq. (12b), whose error will increase with |*x* − *x*_0_|. Nevertheless, despite the visible error under large |*x* −*x*_0_|, the ground truth model evidence *p*(*x*) has little mass under large |*x* − *x*_0_| (Eq. 9, Fig. 2D), and therefore the KL divergence from the true model evidence *p*(*x*) to the circuit’s estimate is still small and acceptable (Fig. 2E).

We emphasize that the circuit model with **fixed synaptic weights** can flexibly compute the posteriors (Fig. 1H-I) and the model evidence (Fig. 2E) **under different parameters** of the likelihood and prior, which are manipulated by changing the location and strength of feedforward and prior input (Eq. 5). The circuits can compute the model evidence sufficiently well when its variance 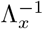 is not too large, which is the regime that the first-order Taylor expansion works well (Fig. 2D-E).

### 5.2 Non-equilibrium circuit sampling: estimates the model evidence lower bound

The above analysis focuses on the circuit sampling in the equilibrium state, e.g., the expectation is under the equilibrium posterior *p*(*z*|*x*) in Eq. (16). Now we consider the non-equilibrium posterior sampling in the circuit (Fig. 3), where the bump position dynamics (Eq. 12a) have not reached equilibrium (so-called burn-in period in MCMC). We denote the *q*_*t*_(*z*_*E*_|*x*) as the distribution of the bump position *z*_*E*_ at time *t* (Eq. 12a), which will gradually converge to the posterior *p*(*z*_*E*_|*x*) in the equilibrium (*t* → ∞). The *instantaneous* decrement of the bump height Δ*Ū*_*E*_ (*x, t*) = *Ū*_*E*_ (*x, t*) − *Ū*_*E*_ (*x*_0_) characterizes the instantaneous logarithm of model evidence ln 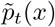 in the circuit, which can be calculated as (Appendix E.1),

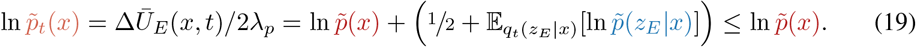

**Figure 3:**
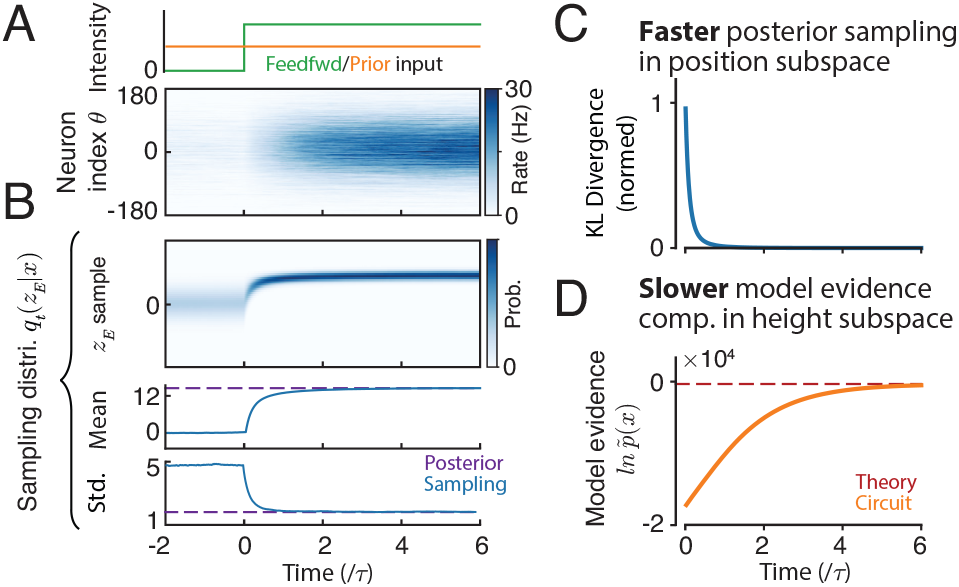
The non-equilibrium model evidence computation in the circuit when just receiving the feedforward input. (A) The transient E population responses. (B) The time evolution of the sampling distribution *q*_*t*_(*z*_*E*_ |*x*). From top to bottom: the sampling distribution density, and its mean and variance. (C) The KL divergence from the true posterior *p*(*z*_*E*_|*x*) to *q*_*t*_(*z*_*E*_|*x*) over time. (D) The circuit’s non-equilibrium estimation ln 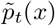.

It provides two insights into the circuit’s non-equilibrium model evidence computation.

1. **Unbiased at the equilibrium**. In the Gaussian case, 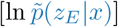 is maximized with maximum of −1*/*2 when *q*_*t*_ = *p* at the equilibrium, suggesting that ln 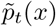 will reach the true model evidence ln 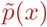 at the equilibrium, i.e., ln 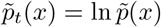 when *q*_*t*_(*z*_*E*_|*x*) = *p*(*z*_*E*_|*x*).
2. **Downward biased at the non-equilibrium**. 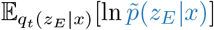 increases *monotonically* when *q*_*t*_(*z*_*E*_ *x*) converges to *p*(*z*_*E*_ *x*), suggesting ln 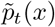 increases monotonically and converges to ln 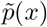 from the lower bound.

Combined, the circuit’s non-equilibrium model evidence estimation ln 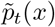 is a lower bound of the true ln 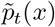, which is analogous to the evidence lower bound (ELBO) in variational inference [29, 57] (Appendix E.1). Nevertheless, a subtle difference is that ELBO deals with the normalized model evidence ln *p*(*x*), while the circuit estimates the unnormalized ln 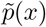 (see Discussions about normalized ln *p*(*x*) calculation in circuits).

The network simulation confirms our theoretical analysis: as the sampling distribution *q*_*t*_(*z*_*E*_|*x*) gradually converges to the true posterior *p*(*z*_*E*_|*x*) (Fig. 3B-C), the circuit’s non-equilibrium estimation ln 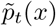 monotonically increases with time and converges to the true ln 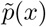 from below (Fig. 3D).

#### Multi-timescale computation in the circuit subspaces

Comparing the Fig. 3C and D, there are two time scales in the circuit subspace computation: the **faster** timescale of the posterior sampling in the bump position subspace (Fig. 3C), and the **slower** timescale of the model evidence estimation in the bump height subspace (Fig. 3D). Eq. (19) provides the explanation for the two timescales: even if the sampling distribution *q*_*t*_(*z*_*E*_|*x*) converges to the posterior *p*(*z*_*E*_|*x*), the expectation in model evidence estimation, i.e., 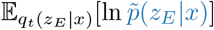, is realized by averaging over stimulus samples effectively collected over time. That is why the model evidence estimation is slower than the posterior sampling.

## 6 Model evidence computation in the circuit receiving multiple sensory inputs

The above analysis only considers the circuit receives one sensory input, whereas the brain often receives multiple sensory inputs that are conditionally independent given the same latent stimulus, e.g., multisensory integration widely studied in neuroscience [4, 58]. We thus extend into the case when the **same circuit** receives multiple feedforward inputs (Fig. 4A). Suppose there are *N* feedforward inputs 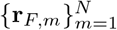 (similar to Eq. 5) independently generated by the same latent stimulus *z* and can have different input strength. Then the stimulus likelihood given all feedforward inputs is,

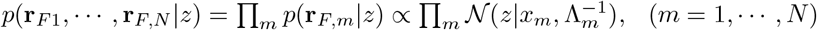

whose mean *x*_*m*_ and precision Λ_*m*_ are similarly determined as (Eq. 6). In this case, the stimulus posterior *p*(*z*|**x** = (*x*_1_, · · ·, *x*_*N*_)^⊤^) and the model evidence *p*(**x**) are,

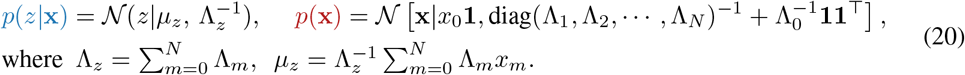

where **1** is a N-dim. vector with all elements being 1. We redo the theoretical analysis on how the same circuit receiving multiple feedforward inputs can compute the posterior and model evidence concurrently in separate subspaces (Appendix F), where the overall analysis rationale is similar to the analysis in one feedforward input case (Sec 4-5). Overall, the theoretical analysis and the network simulation both confirm the **same circuit** can robustly sample posterior and compute multivariate model evidence in response to multiple inputs (Fig. 4B-F). The circuit’s error of model evidence estimate increases with the max eigenvalue of the model evidence’s covariance matrix (Fig. 4D,F).

**Figure 4:**
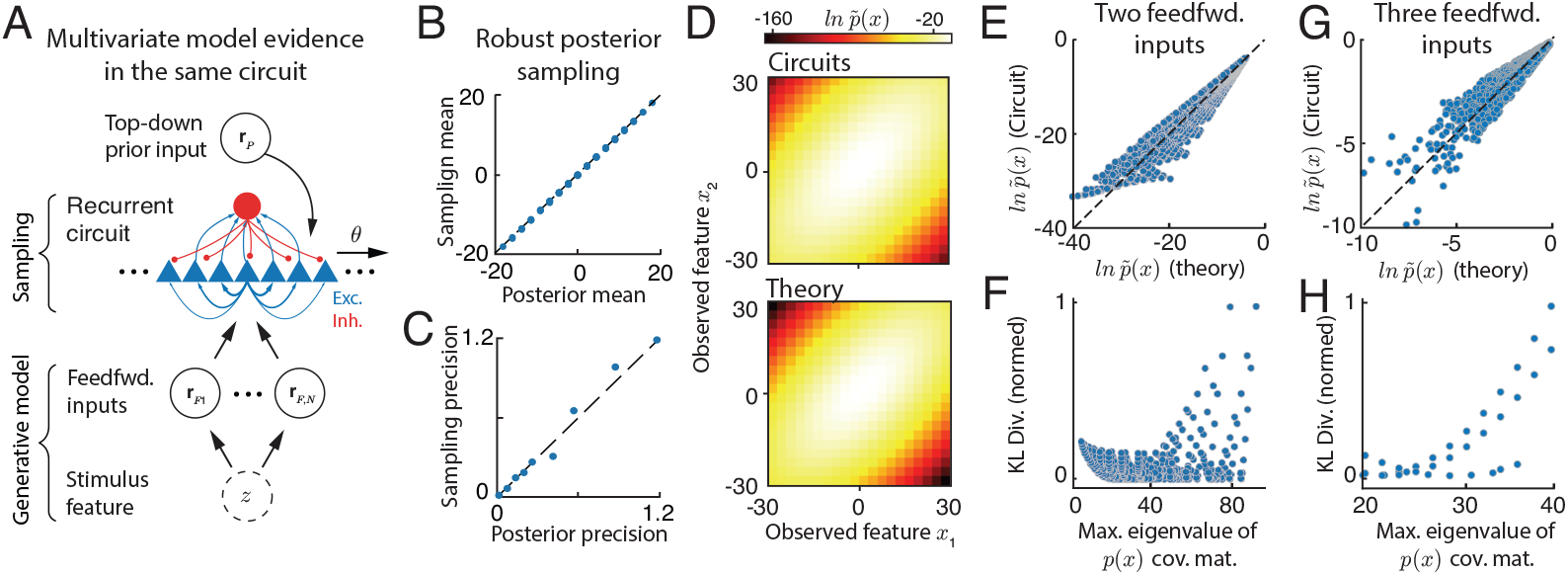
The same recurrent circuit automatically computes the **multivariate** model evidence whose dimension is determined by the number of feedforward inputs. (A) The same circuit receives multiple feedforward inputs. (B-C) The same circuit with fixed weights flexibly samples posteriors with different means and precisions by receiving two feedforward inputs. (D) The logarithm of model evidence ln *p*(*x*_1_, *x*_2_) computed by the circuit (top) and the ground truth (bottom). (E-F) The comparison of the circuit computed ln *p*(*x*_1_, *x*_2_) and the ground truth. Each dot is a result under a combination of likelihood and prior precisions. The circuit’s error, measured by the KL divergence from the ground truth *p*(*x*_1_, *x*_2_) to the circuit’s estimate, increases with the max eigenvalue of the model evidence covariance matrix (F). (G-H) The same as (E-F), but the circuit receives three feedforward inputs and the model evidence becomes *p*(*x*_1_, *x*_2_, *x*_3_).

## 7 Conclusion and Discussion

How the canonical recurrent circuits compute the posterior and model evidence is a fundamental question in computational neuroscience, but it remains largely unknown especially the model evidence computation in the circuits. The present theoretical study discovers for the first time that the recurrent circuit model of CANs can simultaneously realize the posterior sampling and model evidence computation in separate dominant subspaces. The *non-equilibirum* circuit dynamics estimates the model evidence convergently from below, which is analogous to the evidence lower bound in stochastic variational inference. Our study significantly deepens our understanding of Bayesian computation in recurrent neural circuits, revealing the intricate computational structure between circuit subspaces. Our circuit model also has the potential to inspire the network building block for generative modeling in ML, e.g., the latent space sampler.

### Comparison with previous studies

To the best of our knowledge, there is only one previous study investigating the model evidence computation in biological neural circuits ([10]; reviewed by [2]). Rather than considering circuit sampling as in the present study, the earlier study considered a deterministic circuit to implement the marginalization of Gaussian distributions in the model evidence calculation. And it found that the harmonic mean of the precisions during the model evidence computation (Eq. 9) can be computed by a nonlinear transformation whose form is similar to the divisive normalization. Due to the large difference between the circuit dynamics and algorithms in the two studies, it is hard to directly compare them. Considering the flexibility of sampling in approximating arbitrary distributions, the circuit model in the present study may be more scalable to more complex generative models beyond the Gaussian case, which is worth exploring in the future.

### Limitations and future directions

**First**, the present study only considers the posterior and model evidence computation under an embedded linear Gaussian model in subspaces. Note that we don’t manually specify the Gaussian distributions but it naturally emerges from the Gaussian-profile recurrent connectivity (Eq. 3) and input tunings (Eq. 5), which are widely used in recurrent circuit modeling of computational neuroscience. We emphasize even if the Gaussian model is simplified in computation, how in this case the model evidence is computed in the recurrent circuit is still unknown. In the future, it is worth extending to non-Gaussian cases in circuits by using various ways:

i. Change the profile of input tunings and recurrent connectivity to enable the recurrent circuits to compute other distributions belonging to the linear exponential family [9, 19];
ii. Connect the present Gaussian sampling circuit with encoder and decoder to transform the latent Gaussian distributions into more complex distributions. And many deep generative models also consider latent Gaussian sampling, e.g., VAE [29] and latent diffusion model [59]. In this way, our recurrent circuit acts as the <u>latent space sampler</u> in the deep generative models, and potentially seamlessly embed deep generative models into neural networks, rather than merely using neural network as a function approximator to map between input and latent spaces. **Second**, the prior was modeled as a top-down input to the recurrent circuit in the present study, which is a common approach in many network modeling [60, 61]. In contrast, another possibility is that the prior is internally stored in the recurrent connectivity within the circuit through various ways [62, 63], without receiving any top-down input. **Third**, we only consider the univariate latent stimulus, even if the circuit can receive multiple sensory inputs (Sec. 6). To extend to the multivariate latent stimulus inference, we can consider a coupled network model with multiple recurrent circuits [19, 64], where each circuit receives one sensory input and samples the corresponding univariate latent stimulus while the whole coupled circuits can sample the multivariate latent stimulus. It is important to study how the model evidence is computed distributively in the coupled circuits. **Fourth**, our circuit can only compute the (unnormalized) model evidence, ln 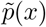. It remains open about whether and how neural circuits compute the normalized evidence *p*(*x*), which is also a hard problem in ML. One possibility is the neural circuits don’t compute the normalized evidence *p*(*x*) but compare each unnormalized evidence 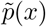 with a reference value to make decisions [5, 11]. All of these form our future research.

## Acknowledgments and Disclosure of Funding

W.H.Z. is supported by the UT Southwestern Endowed Scholars program. The authors thank Libo Ma, Gengshuo Tian, Cheng Xue for fruitful comments on the manuscript.

## Appendix

### A Appendix Figures

**Figure A1:**
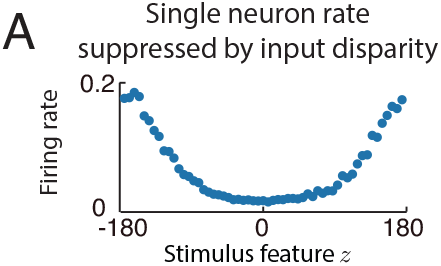
(A) Tuning curve of an excitatory neuron with preferred feature 0°, measured while the sensory likelihood input is moved across the feature space from −180° to 180°.

**Figure A2:**
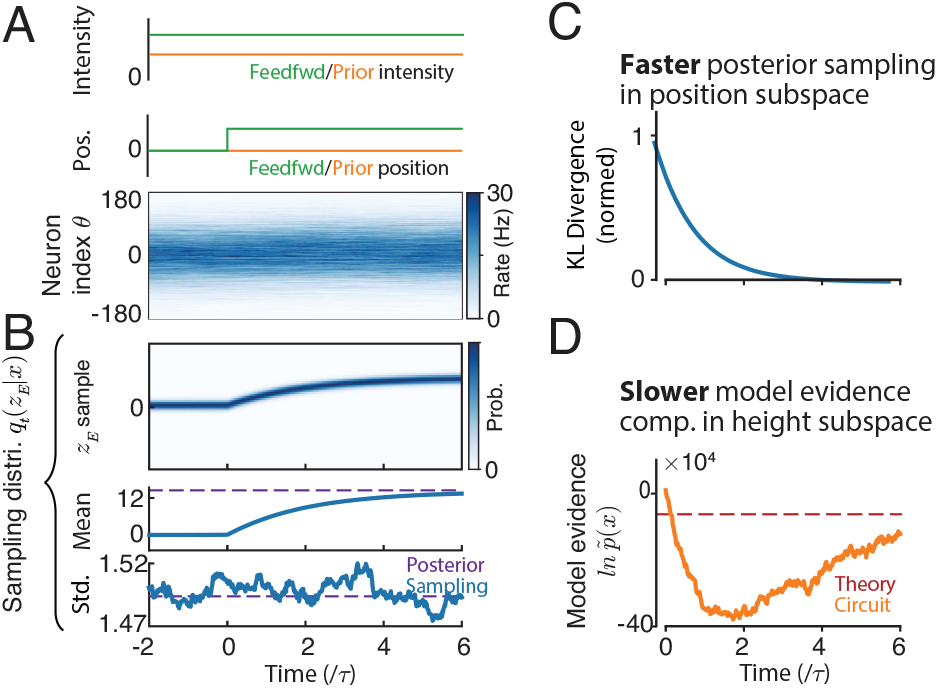
Non-equilibrium circuit dynamics when the likelihood input is abruptly shifted from 0° to 60°, while the prior input location is kept fixed. (B) Transient excitatory population responses. Top: temporal profiles of the prior and sensory feedforward input intensities. (C) Time evolution of the sampling distribution *q*_*t*_(*z*_*E*_ | *x*). From top to bottom: sampling distribution density, mean, and standard deviation. The dashed purple line indicates the posterior mean. (D) KL divergence from the true posterior *p*(*z*_*E*_ | *x*) to the transient sampling distribution *q*_*t*_(*z*_*E*_ | *x*) over time. (E) Circuit’s non-equilibrium estimate of the unnormalized log model evidence, ln 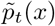, decoded from the bump-height subspace. The dashed red line indicates the theoretical value.

### B Theoretical analysis of the Continuous Attractor Network (CAN)

We present the math of theoretical analyses of the CANs, which is the basis for the subsequent theoretical analyses.

#### B.1 Derivation of the CAN dynamics at the limit of large number of neurons

We start from the conventional spatially discrete form of the recurrent network dynamics,

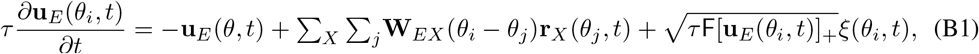

Then we consider the limit of the large number of neurons and the preferred feature of all neurons 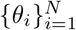 are uniformly distributed in the whole stimulus feature *z* space. And then the neuronal preferred feature effectively becomes a “continuous” value, i.e., *θ*_*j*_ → *θ*. The spatially discrete summation can be effectively converted as

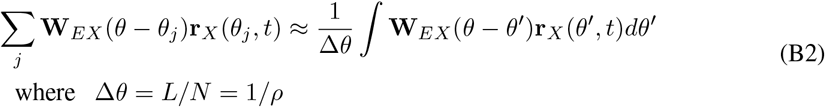

where the Δ*θ* = *L/N* is the interval between the preferred features of two adjacent neurons, which is equal to the length of the stimulus feature space *L* divided by the number of neurons *N* . And thus *ρ* = 1*/*Δ*θ* is the density of neurons in the stimulus feature space. Then the spatially discrete dynamics can be converted into the continuous form as shown in Eq. (2).

#### B.2 Network’s attractor states

We verify the proposed Gaussian ansatz of the attractor states of E neurons (Eq. 10),

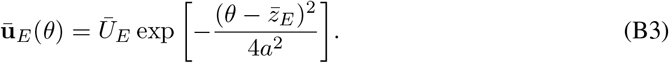

First, we substitute it into the divisive normalization (Eq. 4), yielding the following expression for the firing rate of E neurons,

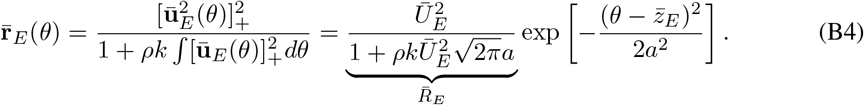

Then we use the above E firing rate (Eq. B4) to calculate the recurrent input from the neuronal population of type *Y* to the one with type *X* in the circuit model,

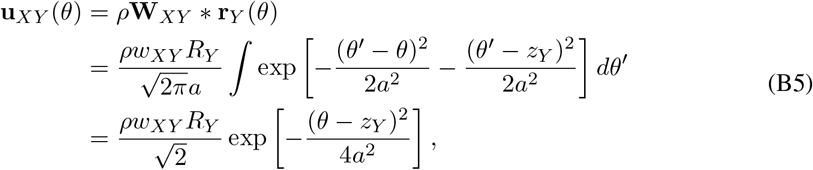

where *z*_*Y*_ is the position of the Gaussian firing rate, and equals to 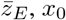 and *x* for recurrent input, prior input, and feedforward input respectively. Specifically, based on Eq. (B5), the three population inputs in Eq. (2) are recurrent E population input is

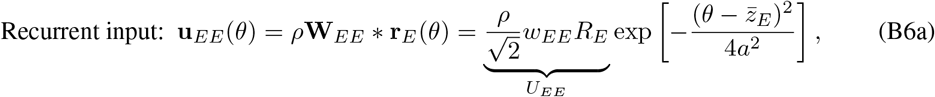

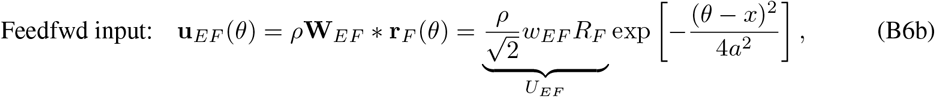

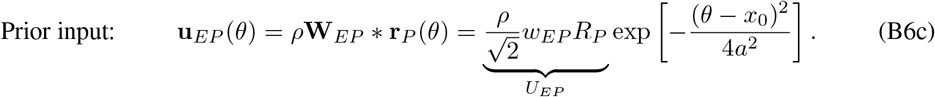

Substituting Eqs. (B3-B4 and B6a-B6c) into the steady state of the E neuron dynamics (Eq. 2), i.e., setting ∂**u**_*E*_*/*∂*t* = 0, we have

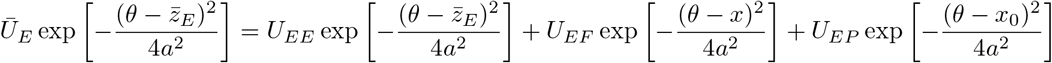

When the feedforward input is located at the same position with the prior input, i.e., *x* = *x*_0_, it can be checked that the E neuron’s position *z*_*E*_ = *x* = *x*_0_ is the only solution to the above equation. And we have,

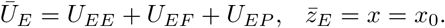

This completes the self-consistency of the Gaussian attractor ansatz (Eq. B3) for the recurrent loop of the dynamics, hence verifies the validity of the Gaussian ansatz.

#### B.3 Dimensionality reduction by projecting on dominant modes

We substitute Eqs. (B3 and B6a-B6c) into the circuit dynamics (Eq. 2). Note that in the temporal dynamics the height and position of E population response (Eq. B3) are time-varying, so we replace the notation *Ū*_*E*_ and 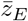 with *U*_*E*_ and *z*_*E*_ respectively, and similarly we replace 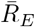 with *R*_*E*_ (Eq. B4), and thus we have

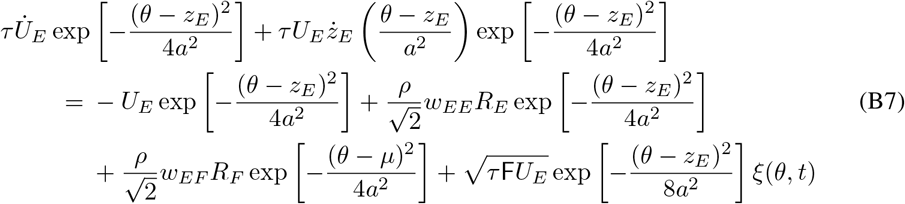

Previous studies analytically calculated the first two dominant eigenvectors of the perturbed dynamics of the above equation [33, 48],

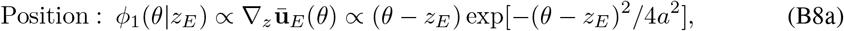

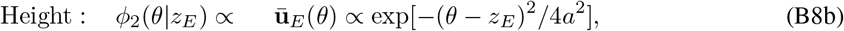

which corresponds to the change of the position *z*_*E*_ and height *U*_*E*_ of the Gaussian bump respectively, Next we project the dynamics Eq. (B7) into these 2 motion modes (Eq. B8). Projection involves computing the inner product ∫ *f* (*θ*)*ϕ*(*θ*|*z*_*E*_)*dθ* with *f* (*θ*) representing every term in Eq. (B7). Projecting onto the bump position model yields the dynamics of bump position *z*_*E*_,

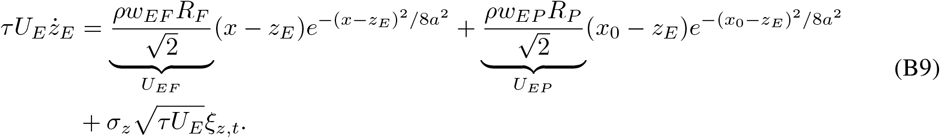

Similarly, projecting onto the bump height model yields the dynamics of bump height *U*_*E*_,

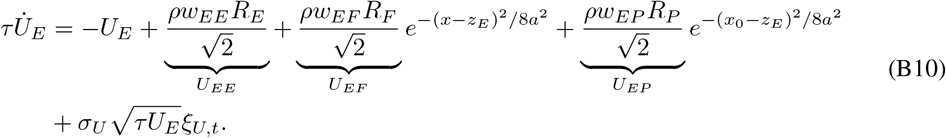

In above equations, the variances of internal variability are,

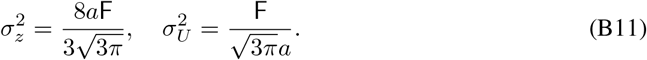

By using the notations of the *U*_*XY*_ in Eqs. (B9-B10) and ignore the Gaussian terms in Eq. (B9), we arrive at the Eqs. (12a-12b) in the main text.

We further explain the rationale behind ignoring the Gaussian terms in Eq. (B9) but keeping the ones in Eq. (B10). This is because the Gaussian term in Eq. (B9) is one order smaller than the ones in Eq. (B10). To see this, we expand the first RHS Gaussian term in Eq. (B9),

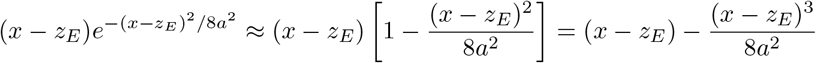

which contains the 3rd order term (*x* − *z*_*E*_)^3^. In comparison, the expansion of the first RHS Gaussian term in Eq. (B10)

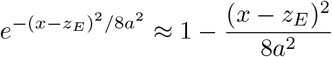

only contains the 2nd order term (*x* − *z*_*E*_)^2^.

### C The probabilistic generative model embedded in the circuit model

#### C.1 The stimulus likelihood

We study how feedforward input defines the latent stimulus likelihood, i.e., ℒ (*z*) ∝ *p*(**r**_*F*_ | *z*). From the Eq. (5), the feedforward input **r**_*F*_ is modeled as a set of independent Poisson spike trains, where each neuron’s firing rate is Gaussian-tuned to the stimulus [2, 5, 9]:

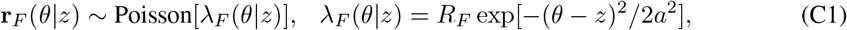

where *λ*_*F*_ (*θ*|*z*) is the mean firing rate of the neuron with stimulus preference *θ*. **r**_*F*_ denotes the peak input rate, and *a* specifies the tuning width. Explicitly writing the Poisson distribution of feedforward input spikes (we discretize the continuous *θ* into equally spaced *θ*_*j*_),

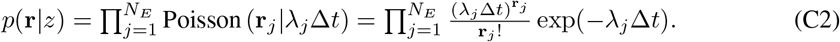

Taking the logarithm,

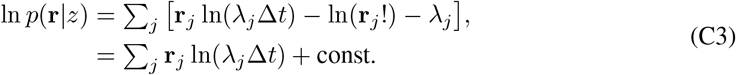

The const. in the above equation is under the assumption that the sum of population firing rate ∑_*j*_ *λ*_*j*_ is a constant irrelevant to latent stimulus *z*, which is true in a homogeneous population with a large number of neurons. Substituting the expression of the Gaussian tuning,

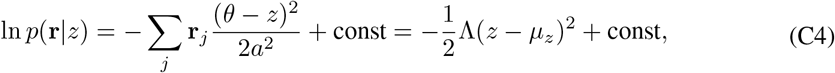

where

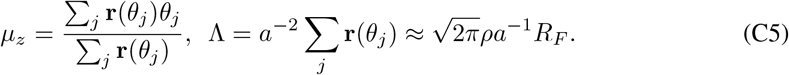

This implies the latent stimulus likelihood for the latent stimulus feature *z* given an observed feedforward input **r**_*F*_ is derived as a Gaussian distribution,

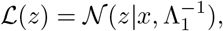

which is the Eq. (6) in the main text. Notably, the Gaussian distribution comes from the profile of the Gaussian tuning (Eq. C1) [9].

#### C.2 The stimulus posterior

Above derivation shows that the feedforward population activity defines a Gaussian likelihood of the latent stimulus feature. Similarly, the prior input centered at *x*_0_ defines a Gaussian stimulus prior,

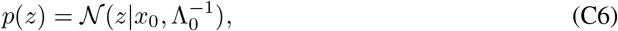

where Λ_0_ is the prior precision. Similar to the likelihood precision in Eq. (C5), when the prior input is implemented by a Gaussian-tuned population with peak rate *R*_*P*_, its precision can be written as

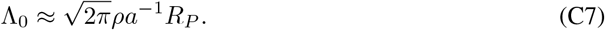

Acrroding to Bayes rule, the posterior distribution can be written as:

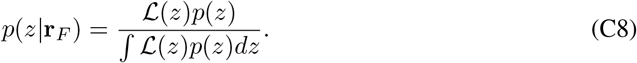

Taking the logarithm of the unnormalized posterior gives

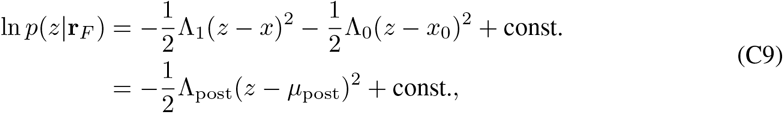

where the posterior precision and posterior mean are

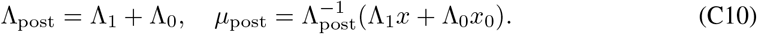

Therefore, the stimulus posterior is also a Gaussian distribution,

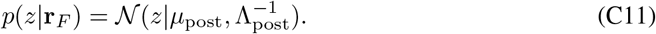

The gradient of the log-posterior is

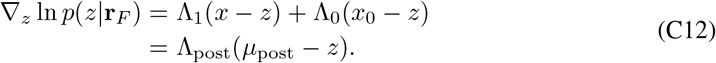

#### C.3 The model evidence

The model evidence is the marginal probability of the feedforward observation after integrating out the latent stimulus feature,

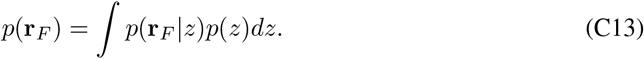

Under the Gaussian approximation above, the likelihood depends on the feedforward population activity through the decoded feature position *x*. Therefore, up to constants independent of *z*, the model evidence can be written as the normalizing constant of the posterior,

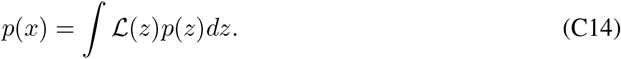

Substituting the Gaussian likelihood and prior gives

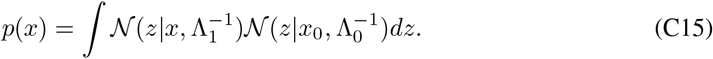

To evaluate this integral, we complete the square,

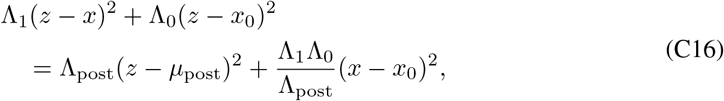

where Λ_post_ and *µ*_post_ are defined in Eq. (C10). Thus the integral over *z* only removes the posterior Gaussian term, leaving

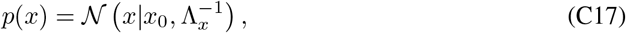

where

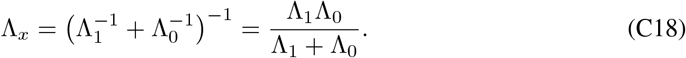

Therefore, the log model evidence is

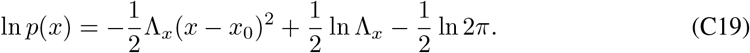

This expression shows that the evidence is high when the feedforward sensory *x* is consistent with the prior center *x*_0_, and decreases quadratically with their mismatch.

### D Langevin posterior sampling in the circuit dynamics

#### D.1 Langevin sampling

The dynamics of Langevin sampling performs stochastic gradient ascent on the manifold of the log-posterior of stimulus features [65], which is written as,

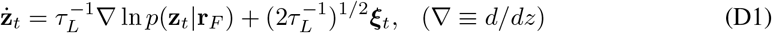

where ***ξ***_*t*_ is a multivariate independent Gaussian-white noise, satisfying 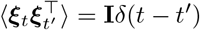, with **I** the identity matrix and *δ*(*t* − *t*^*′*^) the Dirac delta function, and

*τ*_*L*_ is a positive-definite matrix (or a positive scalar in the 1D case) determining the sampling time constant, which is also called the pre-conditioning matrix. Importantly, *τ*_*L*_ is a free parameter of the sampling in that it doesn’t change the equilibrium distribution of **z**_*t*_.

#### D.2 Conditions for realizing Langevin sampling in the circuit

The circuit sampling of the likelihood means the equilibrium distribution of the bump position (Eq. C11) should match with the likelihood (Eq. 6). We copy the circuit bump position dynamics and the likelihood Langevin sampling dynamics in below for comparison,

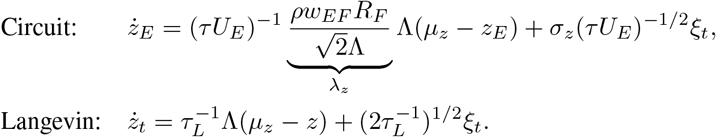

The *σ*_*z*_ is a constant that doesn’t change with neuronal activities. Therefore, the likelihood Langevin sampling in the circuit can be realized by setting the feedforward weight *w*_*EF*_ appropriately to make the ratio of the drift and diffusion coefficients the same as the Langevin sampling dynamics. The optimal feedforward weight can be found as (by using Eq. C5)

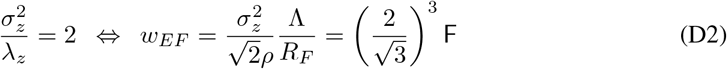

Furthermore, the time constant of the *z*_*E*_ dynamics is

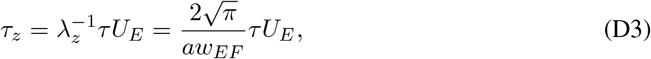

which is proportional to the E bump height *U*_*E*_. Finally, the equation of bump position (Eq. D.2) can be converted into the same form with a standard Langevin sampling,

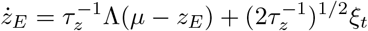

#### E Model evidence estimation by the circuit dynamics and its bias

#### E.1 Downward bias of circuit’s estimation of model evidence

We present the math calculation of the bias of the circuit’s estimation of model evidence, i.e., 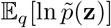 (eq. 19) where *q* is the sampling distribution of the circuit and 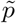 is the unnormalized target density. We show that this quantity is biased upwards and is minimized at *q* = *p*. However, only in the Gaussian case, 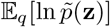 itself is monotonic with respect to *q* and can be used as a surrogate for the ELBO.

Let the instantaneous distribution of bump position in the circuit 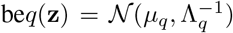 and the unnormalized target:

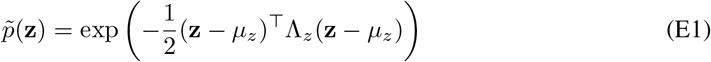

Applying the standard quadratic expectation identity:

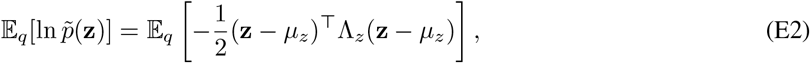

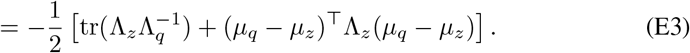

##### Monotonicity analysis

We focus on the monotonicity of 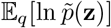 when *q* approaches *p*, which means *µ*_*q*_ → *µ*_*z*_ and Λ_*q*_ → Λ_*z*_. We analyze the two terms in the above equation separately(Fig. A2):

Mean term: (*µ*_*q*_ − *µ*_*z*_)^⊤^Λ_*z*_(*µ*_*q*_ − *µ*_*z*_) ≥ 0, equals zero iff *µ*_*q*_ = *µ*_*z*_. This is the squared Mahalanobis distance, which decreases monotonically as *µ*_*q*_ → *µ*_*z*_.

Variance term: 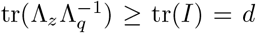 by the trace inequality, with equality iff Λ_*q*_ = Λ_*z*_. Any deviation Λ_*q*_≠ Λ_*z*_ strictly increases the trace.

Therefore 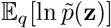 is maximized at *q* = *p*, and increases monotonically as *q* → *p* along both the mean and variance directions.

##### Relationship to normalized density

Since

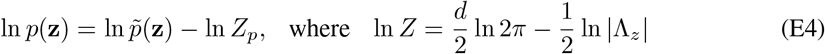

then

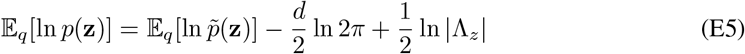

Note that E_*q*_[ln *p*(**z**)] and 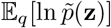 only differs by a constant that is independent of *q*. Therefore, they share identical monotonic behavior as *q* approaches *p*.

##### Connection to ELBO

The evidence lower bound (ELBO) is

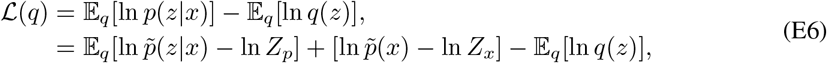

where *Z*_*p*_ and *Z*_*x*_ are the normalizing constants of the posterior and model evidence respectively, i.e.,

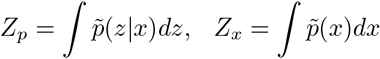

Reorganizing the Eq. (E6),

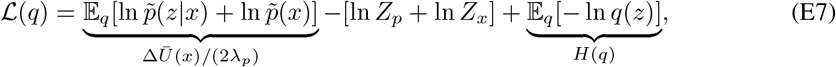

So we obtain the relation between the Δ*Ū* (*x*) and the ELBO ℒ (*q*).

Note that although the ELBO ℒ (*q*) is monotonically increasing when *q* approaches *p*, the entropy term *H*(*q*) is not necessarily monotonic. For example, when *q* is a Gaussian distribution, the entropy is proportional to ln 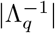, which is not monotonic as Λ_*q*_ → Λ_*z*_. Therefore, the monotonicity of the circuit’s estimation is only a special case under Gaussian case.

#### E.2 Approximate algorithm for model evidence computation

Modern ML has developed sophisticated algorithms to estimate the ln *p*(*x*) via the variational inference, where the ln *p*(*x*) can be estimated via the evidence lower bound (ELBO) ℒ with the help of a variational distribution *q*(*z*),

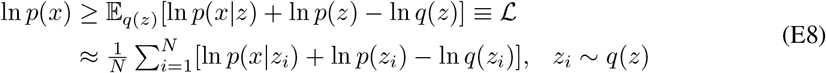

usually the expectation in the ELBO is approximated by Monte Carlo sampling. The gap between the ℒ and ln *p*(*x*) is ln *p*(*x*) − ℒ = *D*_KL_[*q*(*z*) ∥*p*(*z*|*x*), confirming the ℒ is a lower bound of ln *p*(*x*). And the ℒ is tight and equals to ln *p*(*x*) when the *q*(*z*) is exactly the true posterior *p*(*z* | *x*) and thus the KL divergence is zero.

### F. Posterior and model evidence computation: multiple feedforward inputs

#### F.1 The generative model of multiple feedforward inputs

To extend the model evidence calculation to multi sensory case, we fisrt assume, there are *N* feedforward populations 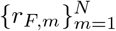, and each of them is conditionally independent given the same latent stimulus feature *z*. Similar to the single-sensory-input case, the *m*-th feedforward population defines a Gaussian likelihood of the latent stimulus feature,

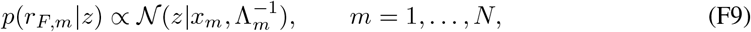

where *x*_*m*_ and Λ_*m*_ are the feature position and precision decoded from sesory input *r*_*F,m*_ by the population vector. The top-down prior input is same as the single sensory input case:

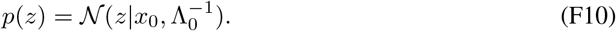

For compact notation, we denote

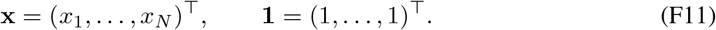

The unnormalized joint density of the latent stimulus and the decoded sensory features is

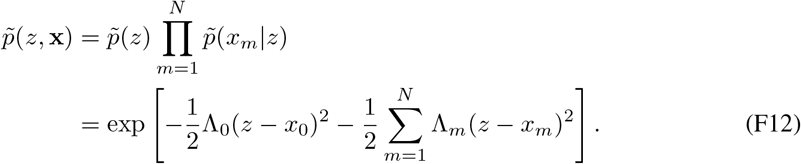

Completing the square gives

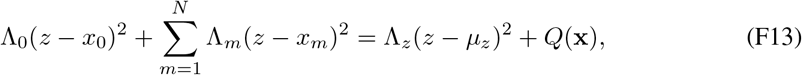

where

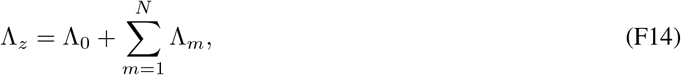

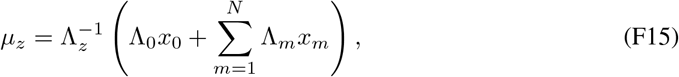

and the residual quadratic term is

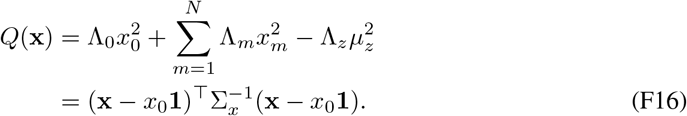

Here the covariance matrix of the multivariate model evidence is

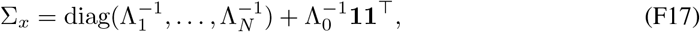

Therefore, the posterior over the shared latent stimulus remains a one-dimensional Gaussian,

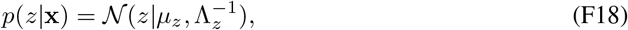

whereas the model evidence becomes an *N* -dimensional Gaussian over the decoded sensory features,

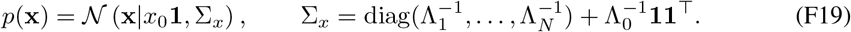

The circuit analysis below concerns the unnormalized model evidence

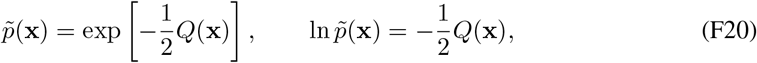

up to a normalization constant independent of **x**.

#### F.2 Bump position dynamics samples the univariate posterior given multi-input

When the circuit receives *N* feedforward inputs, the projected bump-position dynamics becomes

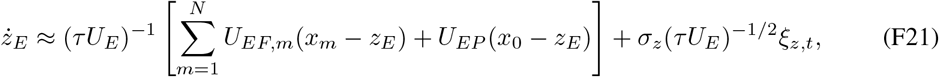

where

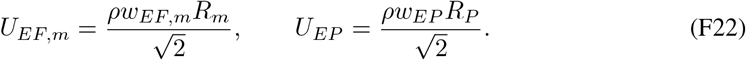

Using the relation between population input magnitude and precision,

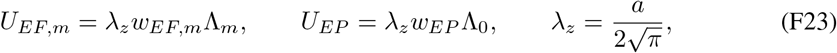

and setting all feedforward and prior weights to the optimal sampling value

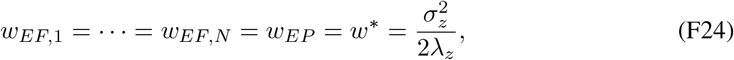

Eq. (F21) becomes

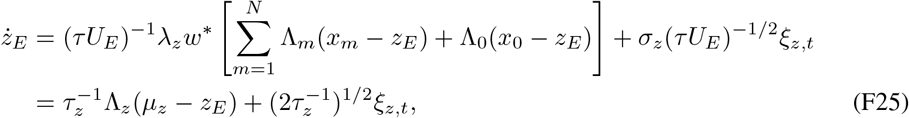

where

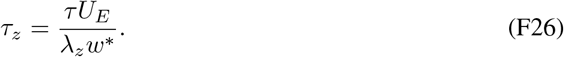

This is the Langevin sampling dynamics whose stationary distribution is 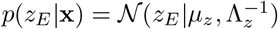. Thus, the same bump-position subspace samples the posterior induced by all sensory inputs.

### G Bump height dynamics computes multivariate model evidence

The projected bump-height dynamics becomes

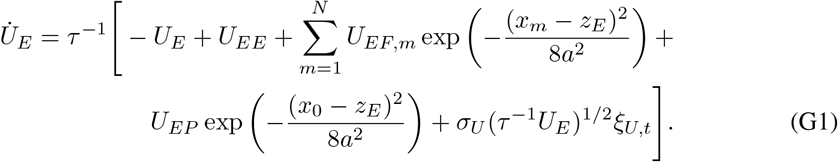

Averaging over the internal noise *ξ*_*U,t*_ and over the equilibrium posterior sampling distribution of *z*_*E*_, and using the first-order Taylor expansion exp[− (*x* − *z*_*E*_)^2^*/*(8*a*^2^)] ≈ 1 − (*x* − *z*_*E*_)^2^*/*(8*a*^2^), we obtain

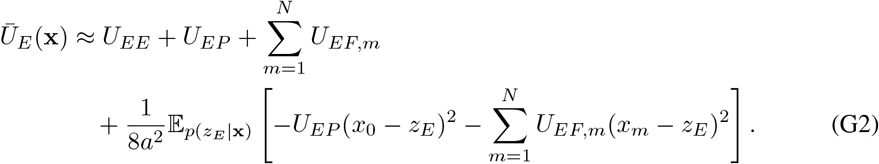

Substituting *U*_*EF,m*_ = *λ*_*z*_*w*^∗^Λ_*m*_ and *U*_*EP*_ = *λ*_*z*_*w*^∗^Λ_0_, and defining

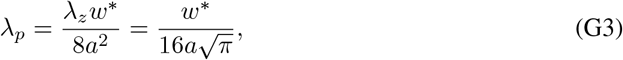

gives

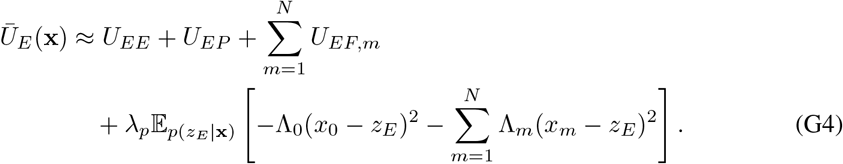

Using the square-completion identity in Eq. (F13),

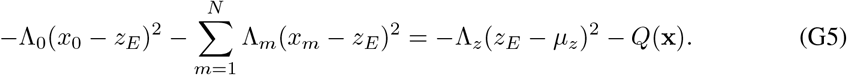

Since 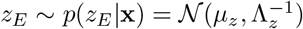 at equilibrium, we have

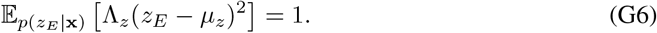

Therefore,

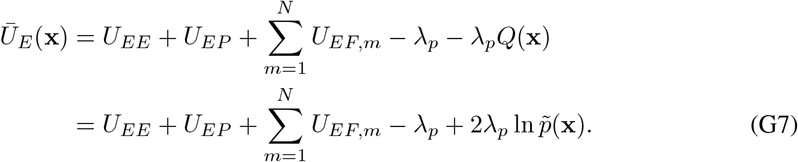

For a fixed set of input precisions, the reference response is obtained when all feedforward inputs are congruent with the prior, i.e., **x** = *x*_0_**1**. In that case *Q*(**x**) = 0 and ln 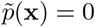, so

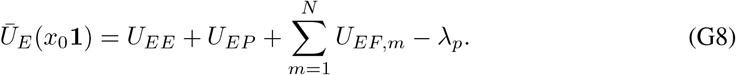

Thus, the decrement of the bump height computes the logarithm of the unnormalized multivariate model evidence:

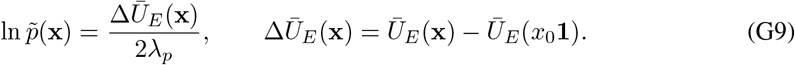

This shows that the same height subspace computes a model evidence whose dimension is determined by the number of sensory feedforward inputs, even though the latent stimulus sampled by the bump-position subspace remains one-dimensional.

### H Circuit simulation parameters and details

#### H.1 Parameters for network simulation

All simulations used the continuous attractor network described in Eq. 2. The network contains one excitatory population whose neurons uniformly tile a one-dimensional circular stimulus feature space, together with a global inhibitory pool implemented through divisive normalization.

The default simulation parameters were shared across all figures and are summarized in Table 1. The stimulus feature space ranged from −180° to 180° and was represented by *N* = 180 excitatory neurons. The neuronal density was therefore *ρ* = *N/*360°. The excitatory time constant was set to *τ* = 1, and the Euler integration time step was Δ*t* = 0.01*τ* . The width of the Gaussian recurrent, feedforward, and prior kernels was set to *a* = 40°. The internal Poisson-like variability was controlled by the Fano factor *F* = 0.5. The strength of divisive normalization was set to *k* = 5 × 10^−4^.

**Table 1:**
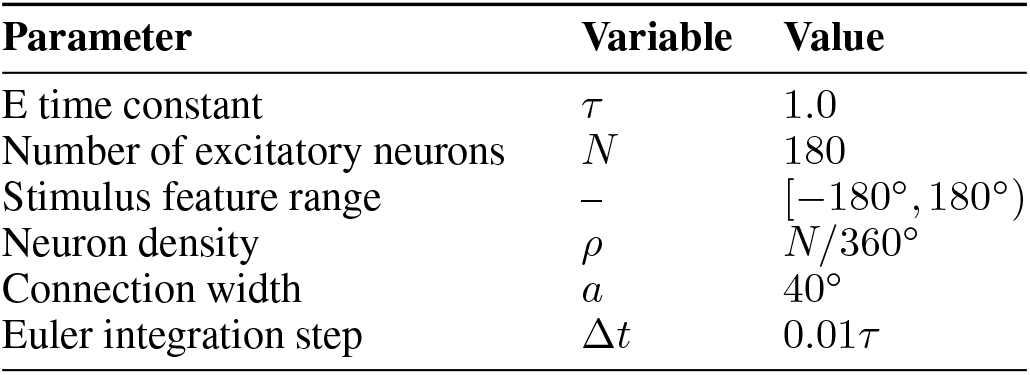
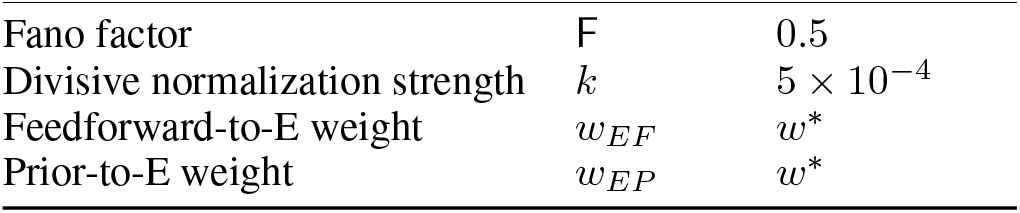
Default parameters used in network simulations.

The feedforward and prior inputs were Gaussian-tuned population inputs of the form

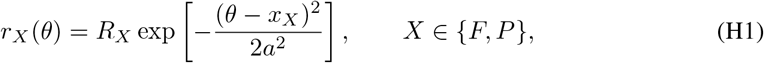

where *R*_*X*_ controls the input magnitude and *x*_*X*_ controls the input location. For the prior input, we denote *x*_*P*_ = *x*_0_ and *R*_*P*_ as the prior center and prior strength. For the feedforward input, *x*_*F*_ = *x* and *R*_*F*_ determine the decoded observation and likelihood precision. In simulations with multiple sensory inputs, each feedforward input *m* had its own location *x*_*m*_ and magnitude *R*_*m*_, while all inputs were projected to the same recurrent circuit.

The feedforward and prior connection weights were set to the theoretically optimal value for Langevin posterior sampling,

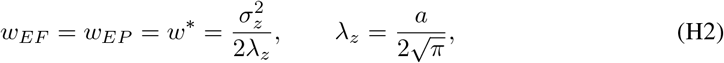

with

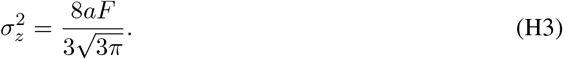

##### Equilibrium simulation

For equilibrium model-evidence simulations, the network was first stimulied with feedforward and prior inputs aligned at the same feature location, to collect the bump height baseline *Ū*_*E*_ (*x*_0_). In rest simulations, the prior location was fixed at *x*_0_ = 0, and the feedforward location *x* was varied across conditions. For each condition, the network activity was simulated for a burn-in period and then recorded for estimating the time-averaged bump height *Ū*_*E*_ (*x*) and the bump-position sampling distribution. The circuit estimate of the unnormalized log model evidence was obtained from the decrement of the bump height,

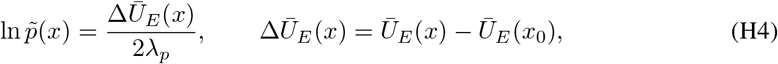

where

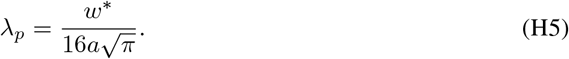

##### Non-equilibrium simulations

For the non-equilibrium simulations, we designed a two-phase protocol to mimic sudden sensory changes. We considered two variants. In the first variant, the circuit received only the prior input during the first phase. At the onset of the second phase, a sensory likelihood input was abruptly introduced at a randomly selected feature location, mimicking the sudden arrival of a sensory stimulus. In the second variant, the prior and likelihood inputs were initially aligned during the first phase. In the second phase, the prior location was held fixed, whereas the likelihood input was abruptly shifted to a new location, mimicking a sudden change in the external environment.The corresponding time-dependent estimate of the unnormalized log model evidence was decoded from the instantaneous bump height,

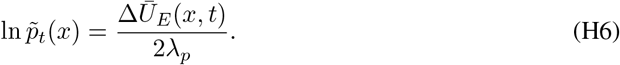

This procedure was used to compare the circuit estimate with the analytical ground truth and to quantify the convergence of the model-evidence estimate.

##### Multi-sensory input simulation

For simulations with multiple feedforward inputs, the same circuit received *N*_*F*_ conditionally independent sensory input populations. We varied the input locations 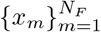 and input strengths 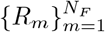 across conditions. The circuit-computed multivariate model evidence was decoded from the same bump-height decrement, using the congruent-input condition 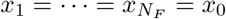 as the reference response.

##### Compute resources

All simulations were run on CPU slurm node with 512G memory. No GPU was required. A single simulation condition for the equilibrium model-evidence experiment took approximately 30s to run.

